# Expression of *Calca* gene-derived peptides in the murine taste system

**DOI:** 10.64898/2026.01.16.700005

**Authors:** Salin Raj Palayyan, Abdul Hamid Siddiqui, Peihua Jiang, Robert F. Margolskee, Sunil Kumar Sukumaran

## Abstract

Taste cell regeneration and taste signaling are regulated by myriad growth factors and signaling molecules secreted by neurons and taste papillae - resident cells. The Calcitonin Related Polypeptide Alpha (*Calca*) gene is a source of four biologically active peptides with varied physiological roles. Alternative splicing of the *Calca* messenger RNA generates either prepro calcitonin gene related peptide (CGRP) or preprocalcitonin encoding transcripts. Proteolytic processing of preprocalcitonin generates procalcitonin, calcitonin and katacalcin. Calcitonin is a ligand for the G-protein coupled receptor calcitonin receptor (CALCR) while CGRP is a ligand for the CGRP receptor (CGRP1R) formed by the calcitonin receptor like receptor (CALCRL)-receptor activity modifying protein 1 (RAMP1) complex. Interestingly, procalcitonin too, is a ligand for the CGRP1R where it can antagonize CGRP. CGRP expression in taste and trigeminal neurons has been documented and is posited to regulate taste signaling. Single cell and bulk RNASeq of taste papillae revealed that the preprocalcitonin but not the *Cgrp* transcript is expressed in *Tas1r3*-expressing type II taste cells, while CGRP1R subunits are expressed in taste stem/progenitor cells and by subsets of fibroblasts and immune cells in the lingual mesenchyme. We confirmed this expression pattern using quantitative polymerase chain reaction (qPCR) and histological techniques. qPCR of geniculate and nodose-petrosal ganglia revealed that both express *Cgrp* and CGRP1R subunit mRNAs, but not preprocalcitonin and *Calcr*. This interesting expression patterns suggests that procalcitonin and CGRP might reciprocally regulate the CGRP1R in the taste papillae and potentially influence taste signaling, taste cell regeneration and the taste microbiome.

## Introduction

Taste cell regeneration, taste neuron pathfinding and taste signaling are regulated by myriad growth factors and signaling molecules.^1,2^ Among the growth factors regulating regeneration, R-spondin 2 is primarily derived from the taste nerve, while sonic hedgehog is derived from both taste nerves and type IV taste cells.^3–7^ Other growth factors such as Wnts, noggin, jagged, bone morphogenetic proteins, insulin like growth factors etc. are presumed to be produced locally by taste and mesenchymal cells in the papillae.^7–11^ Molecules such as nerve growth factor, brain-derived neurotrophic factor, semaphorins and ephrins are required for taste neuron pathfinding, while another set including ghrelin, glucagon like peptide 1, cholecystokinin, neuropeptide Y, leptin and vasoactive intestinal peptide are known to regulate taste signaling.^9,12–21^ The above list is far from exhaustive, and it is likely that many more peptide signaling molecules remain to be discovered. The *Calca* gene is a source of four such molecules (Figure S1A).^22,23^ Tissue specific alternative splicing of *Calca* transcript generates mRNAs coding for either calcitonin gene related peptide (preproCGRP, proteolytically processed to alpha CGRP) or preprocalcitonin (prePCT).^21^ PrePCT is proteolytically processed to generate procalcitonin (PCT), which is further processed to generate calcitonin and katacalcin.^22,23^ All four belong to a larger group of peptides that include beta CGRP, amylin, adrenomedullin, and intermedin, which are ligands for receptors formed by the G protein coupled receptors calcitonin receptor (CALCR) or calcitonin receptor like receptor (CALCRL) by themselves or in combination with one of three receptor activity modifying proteins (RAMP1, RAMP2 or RAMP3).^24,25^ For example, calcitonin is a ligand for CALCR without an associated RAMP subunit, whereas CGRP binds both CALCR+ RAMP1 and CALCRL+RAMP1 (this being its primary receptor, henceforth designated CGRP1R; Figure S1A).^25,26^ CGRP is an exceptionally well-studied neuropeptide; it is expressed in virtually all peripheral sensory neurons, many central neurons, and some non-neuronal cells.^25,27^ It serves a wide variety of functions including nociception, neurogenic inflammation, immunity, vasodilation, wound healing, regulation of the microbiome etc.^27–29^ It is a potent neuro-immune modulator and was recently shown to modulate antigen sampling by M cells in the Peyer’s patch.^27–30^ It displays antimicrobial activity against several gram-negative and gram-positive bacteria and Candida albicans, which adds a further dimension to its immunomodulatory and wound healing roles.^31,32^ From a clinical standpoint, its prominent role in triggering migraine has generated significant scientific and clinical interest. This has led to the development of small molecule- and monoclonal antibody-based CGRP1R antagonists to treat migraines.^27,33,34^ Interestingly, clinical studies have suggested that one-quarter of migraine patients have altered taste perception.^35^ Indeed, CGRP was shown to shape taste transduction by inducing 5-HT secretion by type III taste cells in a phospholipase C-dependent manner.^36^

Calcitonin and katacalcin are key regulators of calcium and phosphate levels in blood and bone.^24,37,38^ PCT is a much less studied neuropeptide. It is not widely secreted in the steady state; it may regulate bone density by suppressing osteoclast (macrophage) migration and maturation.^24,39^ Interestingly, sepsis-associated cytokine storm is preceded by ubiquitous upregulation of PCT expression, leading to its adoption as an early sepsis marker.^40–44^ PCT is a partial agonist of CGRP1R and exerts its mediator role in sepsis through this receptor, and partially antagonizes CGRP at this receptor.^45–47^ However, its biological role(s) in both the steady state and in sepsis remains enigmatic. Analysis of single cell RNASeq (scRNASeq) and bulk RNASeq of the circumvallate papillae (CVP) and fungiform papillae (FFP) showed that the prePCT encoding *Calca*-transcript is highly expressed in *Tas1r3*-expressing type II taste cells in the CVP but only weakly in FFP, while subunits of the CGRP1R are expressed in taste stem cells and lingual mesenchymal fibroblasts and subtypes of immune cells. The CGRP transcript and *Calcr* are not expressed in taste papillae. This was confirmed using qPCR, RNAScope and immunohistochemistry. On the other hand, the nodose-petrosal-jugular (NPJ) and geniculate ganglia that innervate taste papillae were shown to express transcripts for *Cgrp* and CGRP1R subunits, but not the prePCT transcript and *Calcr*. This intriguing expression pattern suggests that taste cell derived PCT and taste nerve derived CGRP may reciprocally modulate CGRP1R signaling in the taste papillae.

## Methods

### Animals

8-10 weeks old C57BL/6J mice (The Jackson Laboratory, Bar Harbor, ME) and Tas1r3-GFP^48^, Lgr5-GFP^49^ and Gad1-GFP^50^ mice strains were used for this study. Animals were housed in a specific pathogen free vivarium with a 12-h light/dark cycle and open access to food and water. All animal experiments were performed in accordance with the National Institutes of Health guidelines for the care and use of animals in research and reviewed and approved by the Institutional Animal Care and Use Committee at University of Nebraska-Lincoln (protocols: 2610 and 2366) and the Monell Chemical Senses Center (protocols: 1163, 1151).

### Bulk RNASeq of taste papillae

10 µm thick coronal sections of fresh frozen tongue from C57BL/6 mice containing FFP or CVP were prepared using a cryostat. Sections were stained with hematoxylin& eosin, and pools of ∼100 taste buds from CVP (n=2 pools from 2 mice) and ∼20-30 from FFP (n=3 pools from 3 mice) were excised using laser micro dissection, and full length bulk RNASeq libraries were prepared using the Ovation RNASeq system V2 (Tecan biosystems, Morgan Hill, CA) per manufacturer instructions. Indexed illumina sequencing libraries were prepared and sequenced in a Hiseq 2000 sequencer (Illumina Inc, San Diego, CA). Raw sequences were aligned to the mouse reference genome (version GRCm38.p3) using the STAR program with default settings and Gencode M14.gtf as the splice junction annotation file, and the reads mapping to genes were counted using the featureCounts package.^51,52^ Normalization and differential expression analysis was done using the DESeq2 package in R/Bioconductor.^53^ Sashimi plots to show alignment of reads were generated using the integrative genome viewer.^54^

### Isolation and scRNASeq of GFP labeled taste cells

scRNASeq of taste cells from Tas1r3-GFP cells from CVP has been previously reported.^55^ scRNASeq of GFP expressing cells from CVP of Gad1-GFP (n=9) and Lgr5 GFP (n=5) mice and FFP of Tas1r3-GFP (n=5) mice was done using the same protocol. Briefly, mice were euthanized by exposure to CO2 followed by cervical dislocation. The tongue was quickly dissected and washed in chilled Tyrode’s solution, and a protease mixture was injected under the lingual epithelium. After a 12 min incubation at 37 °C, the lingual epithelium was peeled, washed, and subsequently incubated in Ca2+-free Tyrode’s for 30 min at room temperature and then replaced with DMEM. Taste buds were triturated with a glass pipette, and individual GFP labeled cells were collected by aspiration using a micro capillary tube controlled by a micromanipulator attached to a fluorescence microscope. aRNA amplification, illumina library preparation and deep sequencing were done as previously described.^55^ Reads alignment, counting, normalization and differential expression analysis was done as described above for bulk RNASeq.

### RNAScope Hiplex assay

RNAscope assay was done using the Hiplex fluorescent assay kit for mice (Advanced Cell Diagnostics Inc., Newark, CA, Cat. no. 324443) with indicated probes (Table S1) as previously described using the manufacturer’s instructions.^56^ Positive and negative control probes were run in parallel to test probes to ensure proper hybridization and imaging conditions were attained in our experiments. Confocal images were captured using a Nikon A1R-Ti2 confocal laser scanning microscope using NIS-Elements A1R software image acquisition and analysis software, using 40/60X objectives. Images were taken using a sequential channel series setting to minimize cross-channel signal, and the channels used were: GFP 488, TxRed 550, Cy5 650 nm. Z-series stack with 10 images per stack was captured at a step size of 1 μm. Acquisition parameters [i.e., gain, offset, photomultiplier tube (PMT) setting] were held constant for experiments. Colocalization counts were made using QuPath software.^57^ Cell boundaries were detected automatically based on DAPI staining. The fidelity of each cell boundary was confirmed by manual inspection. The number of fluorescent spots in each channel (that corresponds to individual mRNA molecules) per cell were extracted and used for colocalization counting. Data from more than two non-consecutive sections from two mice were pooled.

### Immunohistochemistry

Standard immunohistochemical techniques were used as previously described.^56,58^ Briefly, 12 µm thick sections from tongues drop were prepared using a cryostat. Frozen sections were rehydrated with PBS. Nonspecific binding was blocked with SuperBlock Blocking Buffer (Thermo scientific, Waltham, MA, Cat. no. 37580) at room temperature for 1 h. Sections were incubated with primary antibodies overnight at 4 °C in a humidified chamber. After three 15-min washes with PBST, slides were incubated for 1 h at room temperature with the indicated fluorescent secondary antibodies in blocking buffer. All double-immunofluorescent labeling was done with combinations of the secondary antibodies along with DAPI (1:1,000; Invitrogen™, Thermo scientific, Waltham, MA, Cat. no. D1306) to label cell nuclei for cell counting. The primary and secondary antibodies and their concentrations used in this study are listed in Table S2.

Double labeling with two antibodies made from the rabbit (LRMP+PCT and T1R3+PCT), was done as described before.^59^ After blocking, the sections were incubated in succession: first primary anti-rabbit antibody overnight at 4 °C, first fluorescence Fab fragment secondary for 1 h at room temperature, unlabeled anti-rabbit antisera [AffiniPure Fab fragment donkey anti-rabbit IgG (H+L)] for 3 h at room temperature to ensure that all binding sites in the first primary antibody are occupied, second primary anti-rabbit antibody overnight at 4 °C, and then second fluorescence Fab fragment secondary for 1 h at room temperature. Controls for the double immunohistochemistry experiments with two rabbit primary antibodies were done without donkey Fab fragment incubation following the first secondary antibody and omitting the second primary antibody (Figure S4 I & II, control A) or with donkey Fab fragment incubation and omitting the second primary antibody (Figure S4 I & II, Control B) to show adequate blocking of rabbit IgG by donkey Fab fragment. Confocal imaging was done as described above for RNAScope. Colocalization for each taste cell marker was done using images from at least 2 nonconsecutive CVP and FOP containing sections (n=1 mouse). Only those taste cells for which the entire cell body and nucleus could be visualized were counted.

### PCR and qPCR

Total RNA was isolated from freshly dissected geniculate and nodose-petrosal jugular ganglia complex (NPJ, since it is hard to dissect the desired petrosal portion from the NPJ), taste papillae or NT epithelium using the Quick-RNA Microprep kit (Zymo Research Corp, Irvine, CA, Cat. no. R1050) with on-column DNA digestion. cDNAs were prepared from total RNA samples using SuperScript IV VILO Master Mix kit (Thermo Fisher, Waltham, MA, Cat. no. 11756050).

End point PCR and qPCR were done as previously described.^59^ Exon-exon junction spanning primers were designed whenever possible to avoid amplification from any contaminating genomic DNA. A minimum of three biological replicates were used for all cDNA samples. The ratio of the log10 of the average δ-cycle threshold (Ct) value (difference between Ct values of *Bact* and each gene of interest was plotted.^59^ Primers used are shown in Table S3.

## Results

### Bulk RNASeq and PCR show the expression of Calca transcripts and receptors in taste papillae and ganglia

Analysis of bulk RNASeq data from CVP taste buds isolated by laser microdissection showed that the *Calca* transcript is strongly expressed in CVP, but only weakly in the FFP (not shown).

Alignment of the RNASeq reads to the *Calca* genomic locus showed that no reads mapped to the preproCGRP specific exon 5, while large number of reads aligned to prePCT specific exon 4 and to the shared exons 1-3 (Figure 1A). This observation was confirmed using end point PCR using primers specific to the *Cgrp* and prePCT transcripts (Figure S1B), which showed robust amplification for the prePCT transcript in CVP and foliate papillae (FOP) and very weak expression in the FFP. *Cgrp* transcript is expressed at very low levels or were undetectable in all three taste papillae. We could readily detect transcripts for *Calcrl* and *Ramp1* in all three taste papillae and for *Ramp2* in CVP, but not for *Calcr* and *Ramp3*. The *Calcb* transcript that encodes beta CGRP is expressed in all three taste papillae. None of the transcripts are expressed in non-taste lingual epithelium (NT, Figure S1B). Next, we checked the expression of these transcripts in the geniculate ganglia that innervate the FFP and the NPJ ganglia complex, the petrosal portion of which innervate the CVP and FOP. The prePCT transcript and *Calcr* are not expressed in either ganglia, while the *Cgrp*, *Calcrl, Calcb*, *Ramp1*, and *Ramp2* transcripts were detected in both (Figure S2B). Next, we quantified the expression of these transcripts in CVP, FOP, geniculate and NPJ ganglia using qPCR. Consistent with the end point PCR results, robust expression of prPCT is detected in both papillae and weak expression of *Cgrp* is detected in the CVP but not FOP (Figure 1B). Conversely, *Cgrp*, but not prPCT transcript is strongly expressed in both taste ganglia (Figure 1C). Robust expression of *Calcrl, Calcb and Ramp1* were detected in both the taste papillae and ganglia, while *Calcr and Ramp3* are not expressed in any sample. Weak *Ramp2* expression is detected in the two ganglia, while it is not expressed in the taste papillae (Figure 1B-C).

**Figure 1.**
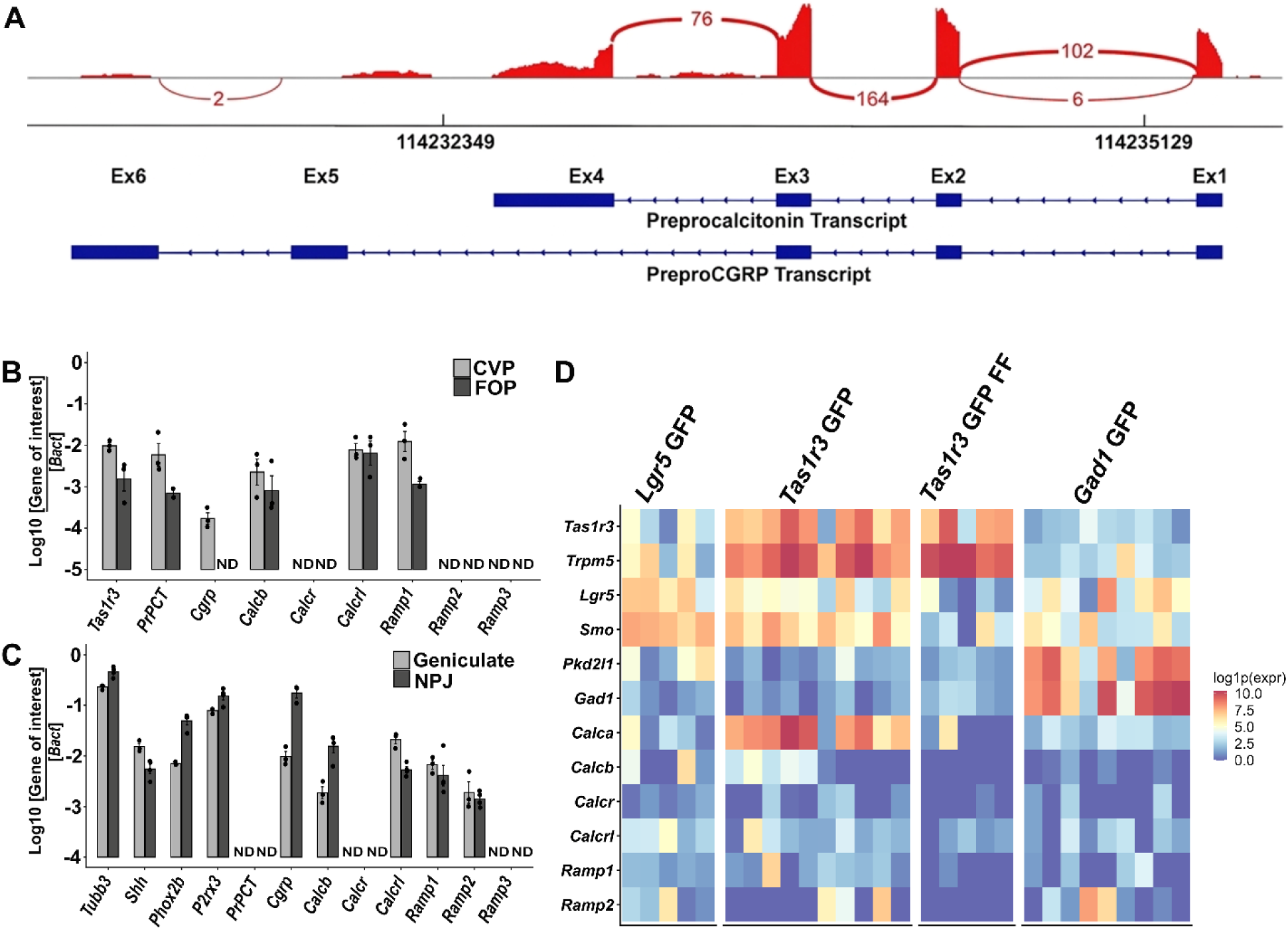
Expression of *Calca* transcripts and receptors for Calca-derived peptides quantified using RNASeq and qPCR. **A)** Sashimi plot of the *Calca* gene locus (transcribed from bottom strand in right to left direction) in a bulk RNASeq library from CVP. The reads aligning to exons 1-6 (Ex1-Ex6, right to left) are shown in red, with the height of the red bars showing the strength of expression, and the number of reads aligning across splice junctions indicated in the loops between exons. Very few reads align to the preproCGRP specific exon 5, while large number of reads align to prePCT specific exon 4 and the shared exons 1-3. **B-C shows qPCR profiling of *Calca and Calcb* transcripts and their receptors in cDNA from taste papillae and sensory ganglia**. **B)** Strong expression of *PrPCT, Calcb*, *Calcrl* and *Ramp1* is observed in both CVP and FOP. Weak expression of *Cgrp* is observed in CVP, while it is undetectable in FOP. *Calcrl and Ramp1* are expressed in both CVP and FOP while *Calcr, Ramp2 and Ramp3 are* not detected in either CVP or FOP. *Tas1r3* is used as a control to demonstrate the quality of taste cDNA. Expression in FFP was not quantified by qPCR due to the preponderance of non-taste epithelium in peeled FFP samples. **C)** qPCR of above transcripts in geniculate and NPJ ganglia. *Cgrp* and *Calcb* are expressed in both ganglia, with stronger expression observed in the nodose-petrosal ganglion. *Calcrl, Ramp1 and Ramp2* are expressed in both ganglia and similar levels. Expression of *Ramp3*, *PrPCT* and *Calcr* is not observed in either ganglion. The taste ganglion marker genes *Tubb3, Shh, Phox2b*, and *P2rx3* are used to demonstrate the quality of ganglia cDNA. The expression of each gene is plotted as the logarithm of the ratio between its cycle threshold value and that of the house keeping gene *Bact*. ND= not detected. **D) scRNASeq data showing expression of transcripts for *Calca, Calcb* and their receptors in taste cells.** Data from Lgr5-GFP (n=5 cells), Tas1r3-GFP (n=10), Tas1r3-GFP cells from FFP (n=5) and Gad1-GFP (n=9) cells are shown. Marker genes for type II cells (*Tas1r3, Trpm5*), Lgr5+ cells (*Lgr5, Smo*), and Type III cells (*Pkd2l1, Gad1*) are also shown as controls.

### scRNASeq of taste cells identifies Calca and CGRP1R subunit expressing taste cells

To identify taste cells that express *Calca* and CGRP1R subunits, we turned to scRNAseq dataset from GFP expressing taste cells isolated from CVP of Lgr5-GFP and Gad1-GFP mice and both CVP and FFP of Tas1r3-GFP mice. Robust expression of the *Calca* transcript (alternative splicing info cannot be extracted from 3’ biased scRNASeq data) is observed from Tas1r3-GFP cells (sweet and umami receptor expressing, type II) in CVP but not in Tas1r3-GFP cells in the FFP, and in Lgr5-GFP (stem/progenitor) and Gad1-GFP (Type III) cells from CVP. The CGRP1R subunits *Calcrl* and *Ramp1* are expressed at moderate levels in Lgr5-GFP cells and in Tas1r3-GFP and Gad1-GFP cells (Figure 1 D).

### Histological analyses confirm the expression of Calca, Calcrl and PCT in taste cells

Next, we turned to histological analyses to confirm the cell type specific patterns of *Calca* and *Calcrl*. RNAscope Hiplex analysis of CVP confirmed the scRNASeq results: Using a *Calca* probe capable of detecting both mRNA isoforms, strong coexpression with the sweet taste receptor subunit *Tas1r3* is observed, with 148/164 (90%) of *Tas1r3* expressing cells coexpressing *Calca,* and 148/149 (99%) *Calca* expressing cells coexpressing *Tas1r3* (Figure 2A-A′′). *Calca* is also coexpressed the canonical type II taste cell marker *Trpm5,* with 131/437 (30%) of *Trpm5* expressing cell coexpressing *Calca*, while 131/149 (88%) of *Calca* expressing cells coexpress *Trpm5* (Figure 2 B-B′′). Most type II cells that do not express *Calca* appear to be bitter taste receptor cells, as less than one tenth of *Gnat3* expressing cells (primarily bitter taste cell marker in CVP) expressed *Calca* (Figure 2C-C′′).^60,61^ *Calca* is not expressed in type III taste cells marked by *Ddc*. (Figure 2D-D′′).

**Figure 2.**
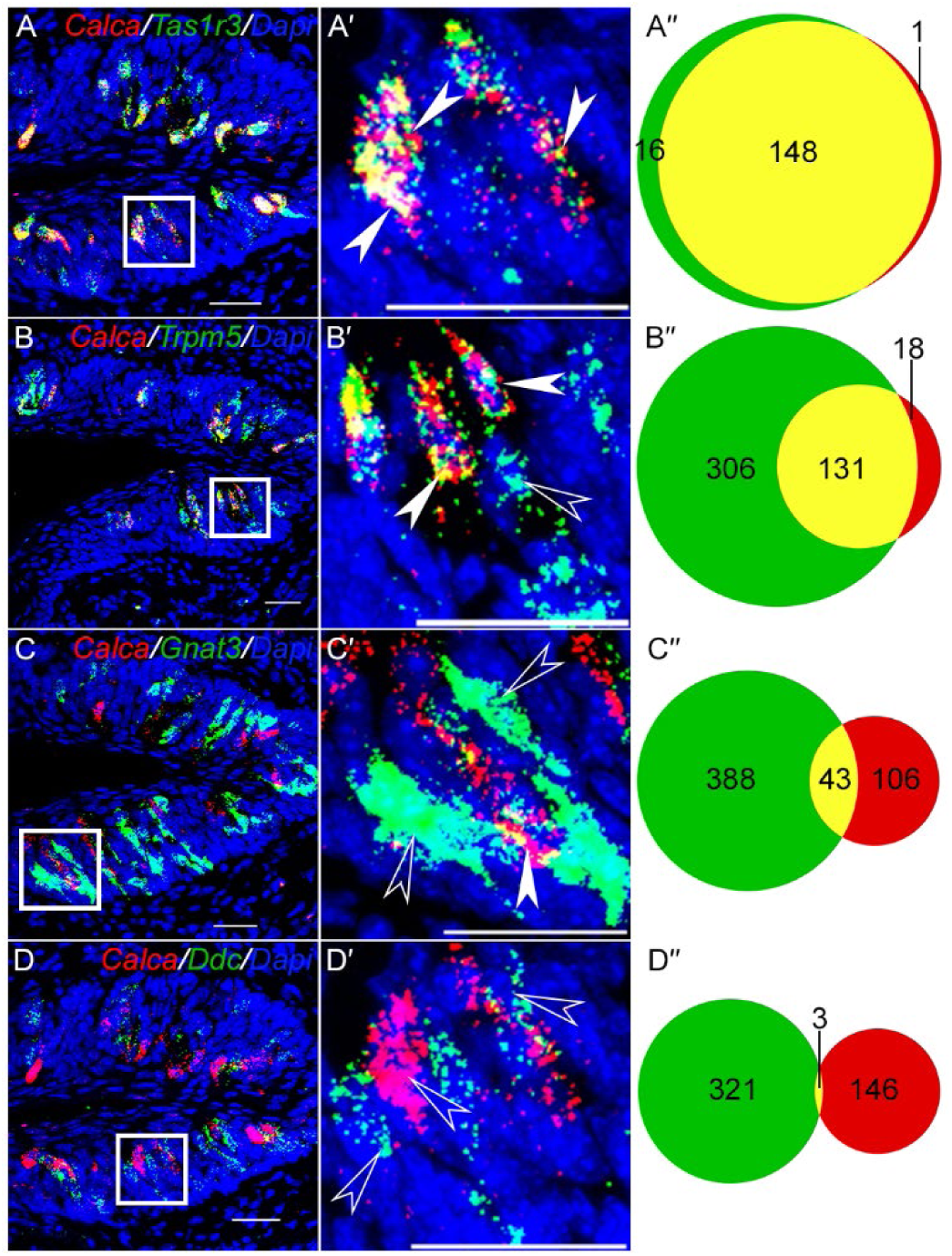
Analysis of *Calca* mRNA expression in CVP. RNAscope Hiplex fluorescence assay was used to determine the coexpression of *Calca* with the taste cell markers *Tas1r3* (A-A′), *Trpm5* (B-B′)*, Gnat3* (C-C′) and *Ddc* (D-D′). The areas highlighted in white boxes in A-D are magnified in A’-D’. Filled white arrowheads highlight double positive cells, and open white arrowheads highlight single positive cells. A′′-D′′ are venn diagrams showing the number of taste cells that co express or singly express the indicated marker genes and *Calca* (Data are from two non-consecutive sections from two mice). Strong coexpression (yellow) of *Calca* is observed with *Tas1r3* and *Trpm5*, less strong coexpression is observed with *Gnat3* and negligible coexpression is observed with *Ddc*. Scale bars = 30 µm. DAPI is used as a counterstain for nuclei.

*Calcrl* staining is observed in unidentified mature taste cells and basal (presumably *Lgr5*+ stem/progenitor) cells in the CVP taste buds (Figure 3A, A′). Significant staining for *Calcrl* is also observed in mesenchymal cells close to basal cells of taste buds that stained strongly for the fibroblast marker *Sparc* (Figure 3A, A′). In agreement with a published scRNASeq dataset from tongue immune cells, *Calcrl* staining was also observed in innate lymphoid cell 3 (ILC3) marker *Ccr6* (Figure 3B, B’) and ILC2 marker *Arg1* (Figure 3 C,C’).^62^ As expected, *Calca* expression is very weak in the FFP, but appear to colocalize primarily with type II taste cells (Figure S2 A-D’). *Calcrl* expression is similar to that in CVP taste buds, with strong expression in mesenchymal fibroblasts and weaker expression in basal and mature cells in the taste buds (Figure S2 E,E’).

**Figure 3.**
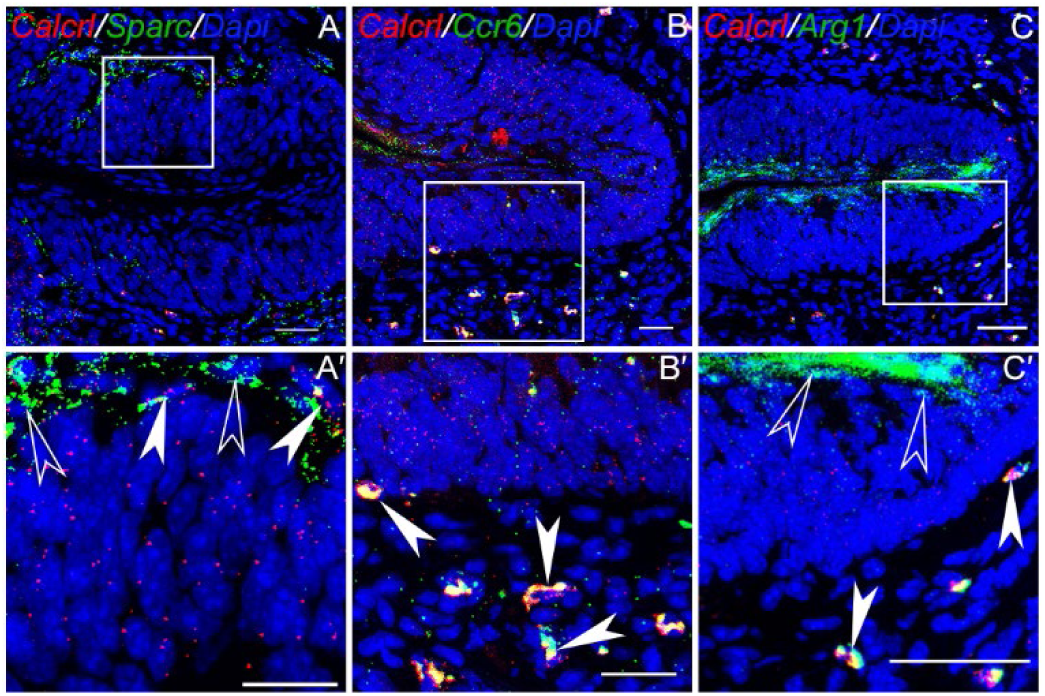
Co expression of *Calcrl* with fibroblast and innate lymphoid cell marker genes. RNAscope Hiplex fluorescence assay was used to determine the coexpression of *Calcrl* with indicated marker genes. Areas inside white boxes in A-C are magnified in A’-C’. A & A′ shows the expression of *Calcrl* in fibroblasts marked by *Sparc* adjacent to basal taste cells. B & B’ shows the expression of *Calcrl* in *Ccr6*+ ILC3 and C & C’ shows its expression in *Arg1*+ ILC2 population of immune cells next to the taste buds. Filled white arrowheads highlight double positive cells, and open white arrowheads highlight single positive cells. Scale bars in A-C are 100 µm and A’-C’ = 30 µm. DAPI is used as a counterstain for nuclei.

Complementary results were obtained using double labeled immunohistochemistry. Using a PCT specific antibody, we saw strong co expression of PCT with the sweet taste receptor subunit T1R3 (100% in both directions) in CVP (Figure 4 A-E) and FOP (Figure 4 F-J). In addition, PCT is strongly coexpressed with the pan-type II marker LRMP, with all PCT expressing cells expressing LRMP and 33/45 (73%) of LRMP expressing cells coexpressing PCT in the CVP (Figure 4 K-O). In case of LRMP, colocalization is observed primarily with cells that stained strongly with LRMP antibody. The weaker LRMP positive cells primarily expressed GNAT3, which, as alluded to before, is primarily expressed in bitter taste receptor cells in the CVP (Figure S3).

**Figure 4.**
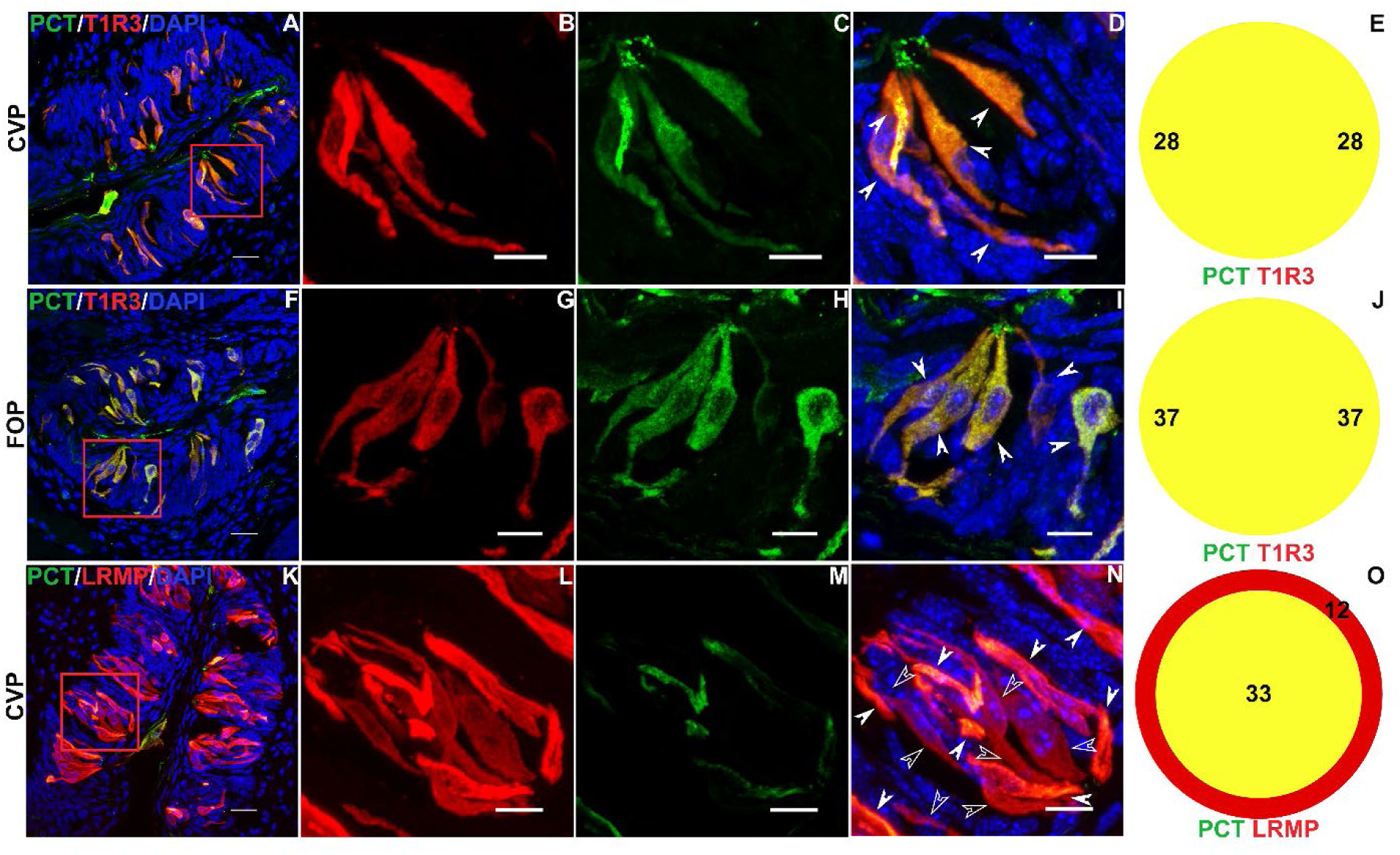
PCT is expressed in type II taste cells. Double labelled immunofluorescence confocal microscopy of CVP and FOP sections with antibodies against PCT (green) and type II taste cell markers T1R3 (red; A–D, F-I) or LRMP (red; K–N). Nuclei are counterstained with DAPI (blue). A, F, K are lower magnification images and dashed red boxes indicate regions shown at higher magnification in respective panels to the right. (D, I, N) Double positive cells are highlighted with solid arrows and single positive cells with hollow arrows. Colocalization counts are shown in Venn diagrams (E,J,O). Scale bars = 20 µm. DAPI is used as a counterstain for nuclei.

### CGRP expression in lingual taste and non taste papillae

Next, we used immunohistochemistry with antibodies against CGRP and TUBB3 to examine the expression of CGRP in neuronal fibers that innervate the lingual epithelium. In agreement with previous reports, strong CGRP expression was observed in fibers that innervate the CVP, with a large proportion of TUBB3 positive nerve fibers coexpressing CGRP.^36^ Processes in the CVP mesenchyme are strongly labeled, as are many fibers that innervated taste buds (Figure 5 A-D). A similar pattern of expression is observed in the FOP and FFP (Figure 5 E-H and I-L).

**Figure 5.**
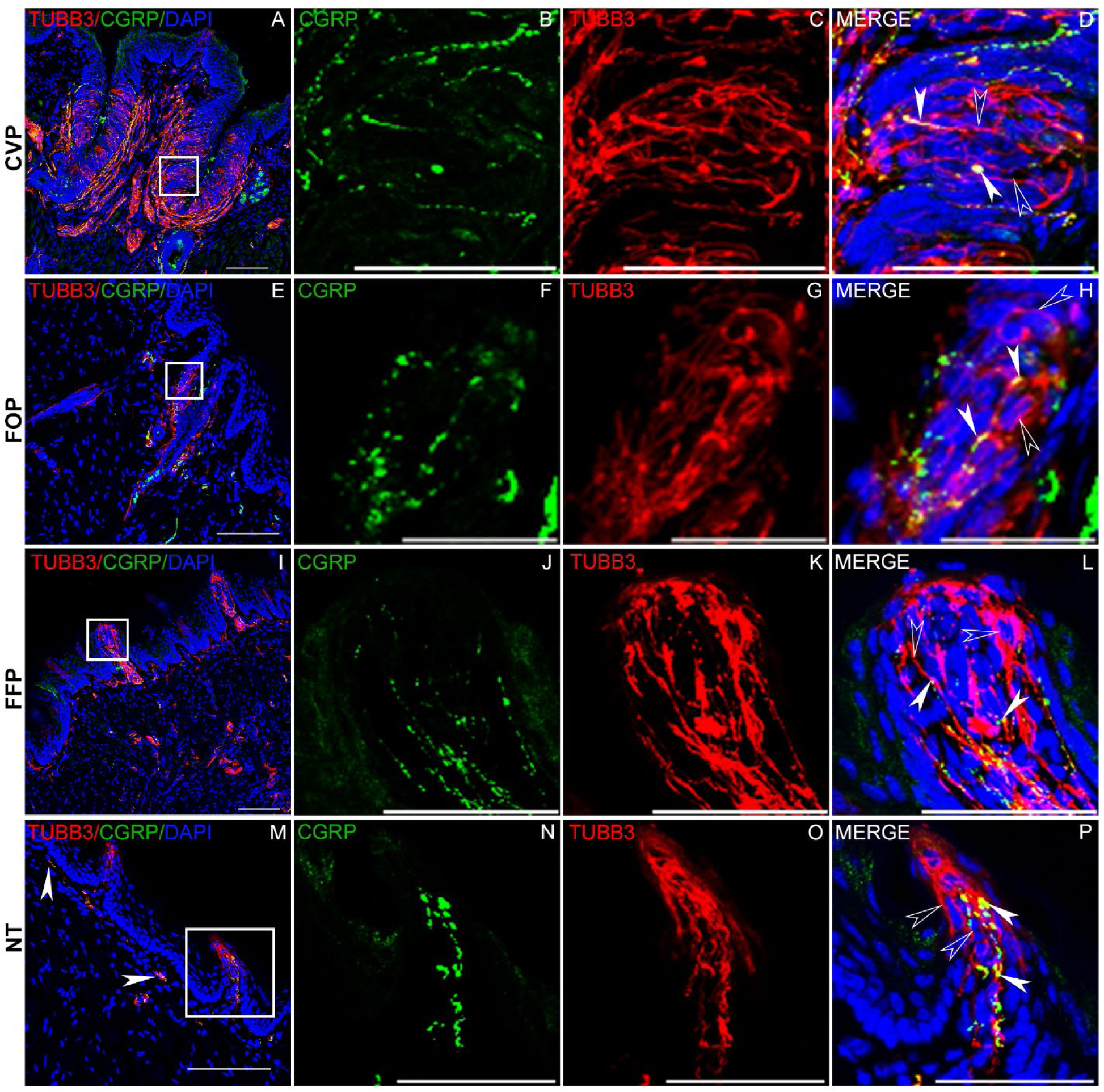
Expression of CGRP in lingual neurons. Double labelled immunofluorescence confocal microscopy with antibodies against CGRP (green) and TUBB3 (pan neuronal marker protein, red) was used to visualize CGRP expression in CVP (A-D), FFP (E-H), FOP (I-L) and non-taste lingual epithelium (NT, M-P). Areas inside white boxes in A, E, I and M are magnified in the respective panels to the right. Strong CGRP expression is seen in a large subset of neuronal fibers in the mesenchyme as well as within taste buds in all three taste papillae. Filled white arrow heads show colocalization, and open white arrowheads show TUBB3 only fibers. Neuronal innervation is weak in the NT epithelium, but the filiform papillae are strongly innervated by neuronal fibers, many of which are CGRP positive. Scale bars in A, E, I and M are 100 µm and those in the remaining panels are 30 µm. DAPI is used as a counterstain for nuclei.

Interestingly, CGRP is also expressed in the fibers innervating the non-taste epithelium, albeit less densely compared to the taste epithelium (Figure 5 M). However, strong CGRP expression is observed in fibers innervating the filiform papillae (Figure 5 N-P). CGRP staining in all parts of the tongue is punctate, likely reflecting localization to vesicles.

## Discussion

Neuropeptides and growth factors are essential for taste function, but it is likely that many key neuropeptides required for taste function have not yet been identified. The *Calca* gene derived peptides CGRP, calcitonin, and procalcitonin play key roles in health and disease.^24,27,33,38–40,63^ Using a number of techniques probing the expression of the corresponding mRNAs and proteins, we show that the prePCT encoding transcript is strongly expressed in CVP and FOP, almost exclusively in the *Tas1r3* expressing type II (primarily sweet and umami receptor expressing) taste cells. Curiously, *Calca* expression (both prePCT and *Cgrp* mRNAs) is very weak in FFP (Figures S1B, S2). The related *Calcb* gene is expressed in all three taste papillae. All three taste papillae express the CGRP1R subunits *Calcrl* and *Ramp1* (Figures 1, 3, S1B, S2). Interestingly, the taste neurons in the geniculate and NPJ ganglia that innervate the papillae expresses the *Cgrp* but not the prePCT transcript and express both the CGRP1R subunits (Figure S1B, Figure 1C). A small amount of *Cgrp* transcript is detected in the CVP using both endpoint and qPCR, although it is likely derived from the nerve endings that innervate the taste papillae rather than taste cells themselves (Figure S1B, Figure 1). This is supported by the absence of *Cgrp* transcript in the bulk RNASeq data from CVP, derived from laser microdissected taste buds devoid of contamination from nerve bundles in the CVP core that would be present in the CVP samples used for PCR experiments (Figure 1A). CGRP expression in trigeminal neurons is well studied, and it is very likely that trigeminal neurons that innervate the taste papillae also express CGRP, although this has not been experimentally demonstrated to our knowledge.^64^ Thus, it is possible that the bulk of alpha CGRP in the taste papillae is nerve derived. Indeed, abundant staining for CGRP is observed in nerves innervating the taste papillae and non-taste lingual epithelium (Figure 5). Previous immunolocalization studies of CGRP in rodent, pig and human taste papillae show CGRP expression restricted to nerve bundles and their termini in and around CVP taste buds.^36,65–67^ PrePCT is processed to generate PCT and then to calcitonin and katacalcin (Figure S1A). We were able to detect the robust expression of prePCT transcript using end point and qPCR (Figures 1 and S1). Although the RNAScope probe for *Calca* does not distinguish between the CGRP and PCT encoding mRNA isoforms, we obtained strong staining in TAS1R3- and LRMP-expressing taste cells with the PCT antibody (Figures 2, S2 and 4). However, these experiments do not allow us to determine if calcitonin and katacalcin are produced by taste cells. Since both peptides are part of PCT, they cannot be readily distinguished using antibody staining. Notably, their biological roles are distinct from that of PCT; both are well known hormones produced by the parathyroid gland that bind to the calcitonin receptor to regulate bone and serum calcium and phosphorus levels.^24,37,38^ PCT on the other hand, is an early marker of sepsis, which is only expressed at very low levels in healthy individuals.^39–42,45^ Notably, we did not detect the calcitonin receptor mRNA (*Calcr*), in the taste papillae or the geniculate and NPJ ganglia using qPCR and only low levels were detected in the scRNASeq data (Figures 1, S1). Thus, the available evidence indicates that calcitonin and katacalcin are either not produced by taste cells and nerves or will not be biologically active in case they are produced. PCT on the other hand, can stimulate CGRP1R, and it also antagonizes CGRP at this receptor.^45,47^ Considering abundant CGRP1R receptor expression in CVP, FOP, and taste ganglia, PCT is the only peptide generated from taste cell expressed prePCT transcript capable of exerting its biological role.

Our model for the likely roles of PCT and CGRP in the taste system is illustrated in Figure 6. There is a large body of literature about the biological roles of CGRP. One study that looked at the effects of CGRP in isolated taste buds using calcium imaging and bioassay showed that CGRP may regulate taste signaling by regulating 5-HT signaling by type III cells.^36^ We (and the original study) did not detect CGRP1R expression in type III cells. We found that *Calcrl* is expressed in taste stem cells using scRNASeq and RNAScope (Figures 1, 3, S2). Moderate *Ramp1* expression in *Lgr5*+ cells is detected in the scRNASeq data, and we could detect it in taste papillae using qPCR as well (Figure 1). CGRP regulates stem cell maintenance, and CGRP1R expression in Lgr5+ cells in CVP and basal (likely stem/progenitor) cells in the FFP taste buds and adjoining fibroblasts raises the possibility that taste nerve derived CGRP regulates taste stem cells directly, and indirectly through the fibroblasts (Figures 1, 3, S2).

**Figure 6.**
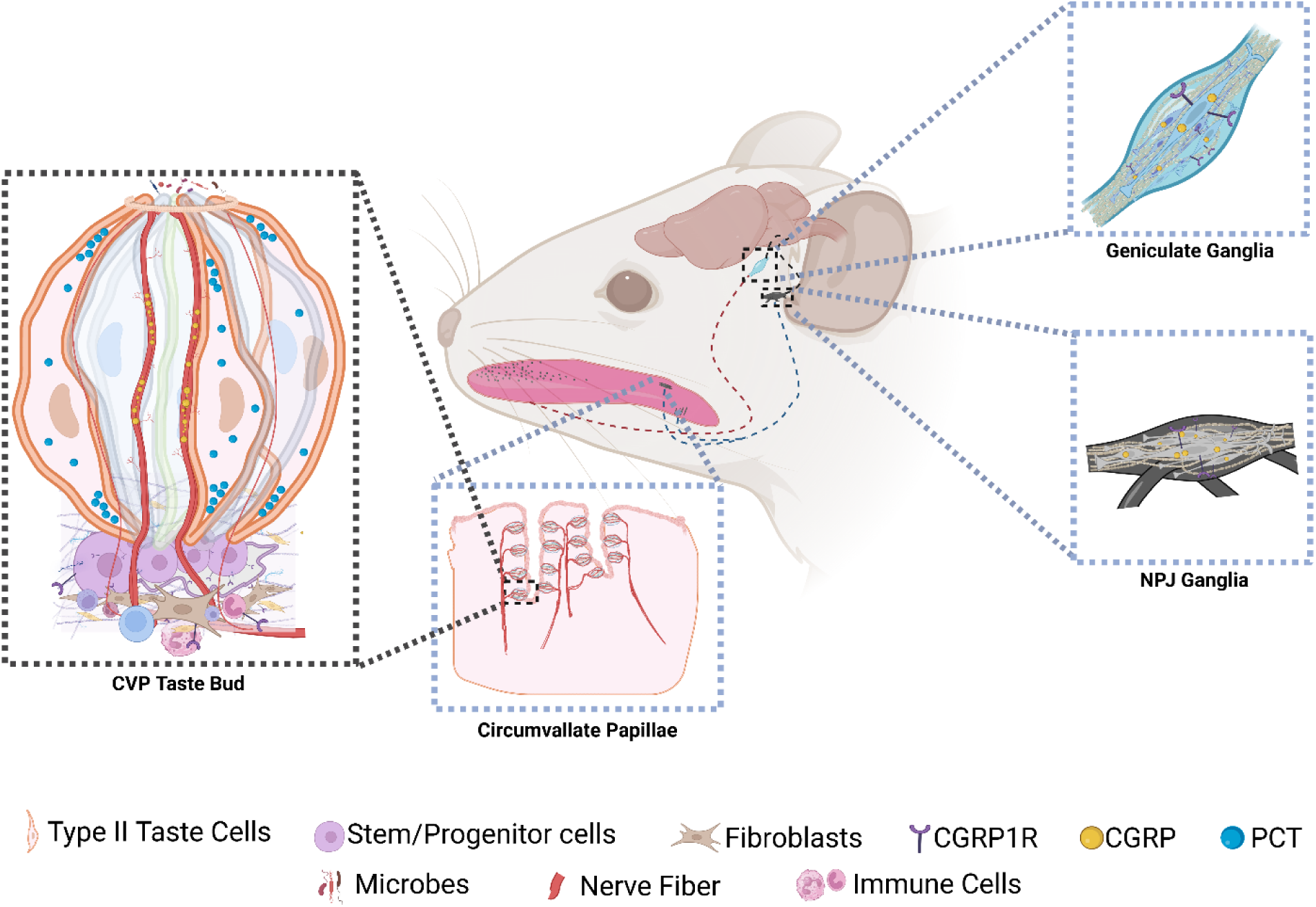
Model for the roles of PCT and CGRP in taste papillae. Lingual CGRP is produced primarily by the taste (geniculate, petrosal portion of NPJ) and trigeminal (not shown) ganglia and delivered to the tongue through the nerve fibers. In CVP and FOP, taste buds are arranged along the walls of trenches that limit their cleansing by salivary flux, likely causing higher bacterial biofilm formation which might proceed to infection if unregulated. PCT produced by type II taste cells in the CVP and FOP, and nerve-derived CGRP can bind to the CGRP1R expressed in some mature taste cells, stem/progenitor cells, mesenchymal fibroblasts and ILC2 and ILC3 cells. They may also be secreted to the taste papillae trenches and modulate the microbiome by virtue of their microbicidal or microbiostatic effects. Illustration created in BioRender. Siddiqui, A. H. (2026) https://BioRender.com/d1kq5qf

CGRP can also regulate immunity at the taste cells through its effects on immune cells, epithelial cells and fibroblasts, and can shape the oral microbiome directly through its microbicidal effects as described above (Figure 6). Intriguingly, CGRP1R is expressed in taste bud-adjacent ILC2 and ILC3 (Figure 3 B-C’). CGRP1R signaling in lungs is a critical regulator of ILC activity, and it plausible that they might play a similar role in taste cells.^68–70^ It is also capable of regulating microfold cells that mediate microbial transcytosis in the Peyer’s patch.^30^ We have shown that type II taste cells might mediate immune surveillance similar to microfold cells, and it is possible that CGRP plays a similar role in taste papillae.^56^ The amount of PCT produced by the taste cells will not be sufficient to elevate it’s circulating levels in the body, and it likely exerts its effects by paracrine signaling within the taste papillae (Figure 6). Sepsis-associated cytokine storm is preceded by ubiquitous PCT expression, leading to its adoption as an early sepsis marker.^40^ However, very little is known about its role in normal physiology. In the skeletal system, it may regulate bone density by suppressing osteoclast (macrophage) migration and maturation.^39^ It’s expression in CVP and FOP but not FFP might provide some clues to its function in taste cells. Unlike the FFP, the CVP and FOP have deep trenches around which the taste buds are arranged. The trenches are relatively less exposed to salivary flux and may be more hospitable to the oral microbiota, including those responsible for halitosis, transient lingual papillitis etc.^71,72^ This raises the possibility that PCT might regulate the taste papillae microbiome in much the same manner as CGRP (Figure 6). In addition, it has the potential to regulate taste signaling and taste cell regeneration by regulating stem/progenitor cell expressed CGRP1R. As stated before, PCT is a partial agonist of the CGRP1R, and it can partially antagonize the effects of CGRP at this receptor. Thus, it is plausible that PCT and CGRP reciprocally shape the biological effects CGRP1R signaling in the taste papillae (Figure 6). In light of this, we hypothesize that the taste papillae might be a suitable model system to determine the biological roles of PCT and its cross talk with CGRP.

## Acknowledgements

We would like to thank Ichiro Matsumoto for the T1R3 antibody. The Microscopy work was carried out at the Microscopy Research Core Facility of the Center for Biotechnology at UNL, which is partially funded by the Nebraska Center for Integrated Biomolecular Communication COBRE grant (P20 GM113126 and NIGMS) and the Nebraska Research Initiative.

## Funding

This work was supported by National Science Foundation CAREER award 2443659, NIH/NIDCR New Investigator RO3 award 1R03DE032417, a Project leader award from Nebraska Center for the Prevention of Obesity Diseases (NIH P30GM154608), and PA State tobacco grant STA019A01SUKUM to SKS and NIH/NIDCD R01DC018627 to PJ.

## Conflict of Interest

The authors declare no conflict of interest.

## Author Contributions

SRP and AHS: performing experiments, acquiring the data, data analysis, interpretation of the results, creation of the figures, and revision of the manuscript. PJ and RFM: funding acquisition, supervision of data acquisition, interpretation of results and revision of the manuscript. SKS: funding acquisition, conceptualization and design of the study, supervision of data acquisition, interpretation of results, writing of the manuscript.

## Data availability

The underlying bulk and scRNASeq data are being uploaded to NCBI’s short read archive and will be available very soon.

## Notes

### Competing Interest Statement

The authors have declared no competing interest.

### Summary of Updates

We have added new data on the coexpression of Calca with innate lymphoid cell (ILC2 and ILC3) marker genes, coexpression of Calca and Calcrl with taste and fibroblast marker genes in the fungiform papillae, and of CGRP in neurons innervating all four lingual papillae and the non-taste epithelium. We realize that the droplet based scRNASeq dataset used in the initial submission requires a standalone publication to fully describe it, which is under preparation. Hence, we have replaced it with an earlier set of scRNASeq data derived from GFP labeled taste cells generated in collaboration with Drs. Margolskee and Jiang, who are now included as co-authors.

