## Supplementary Data for "Expression of *Calca* gene-derived peptides in the murine taste system"

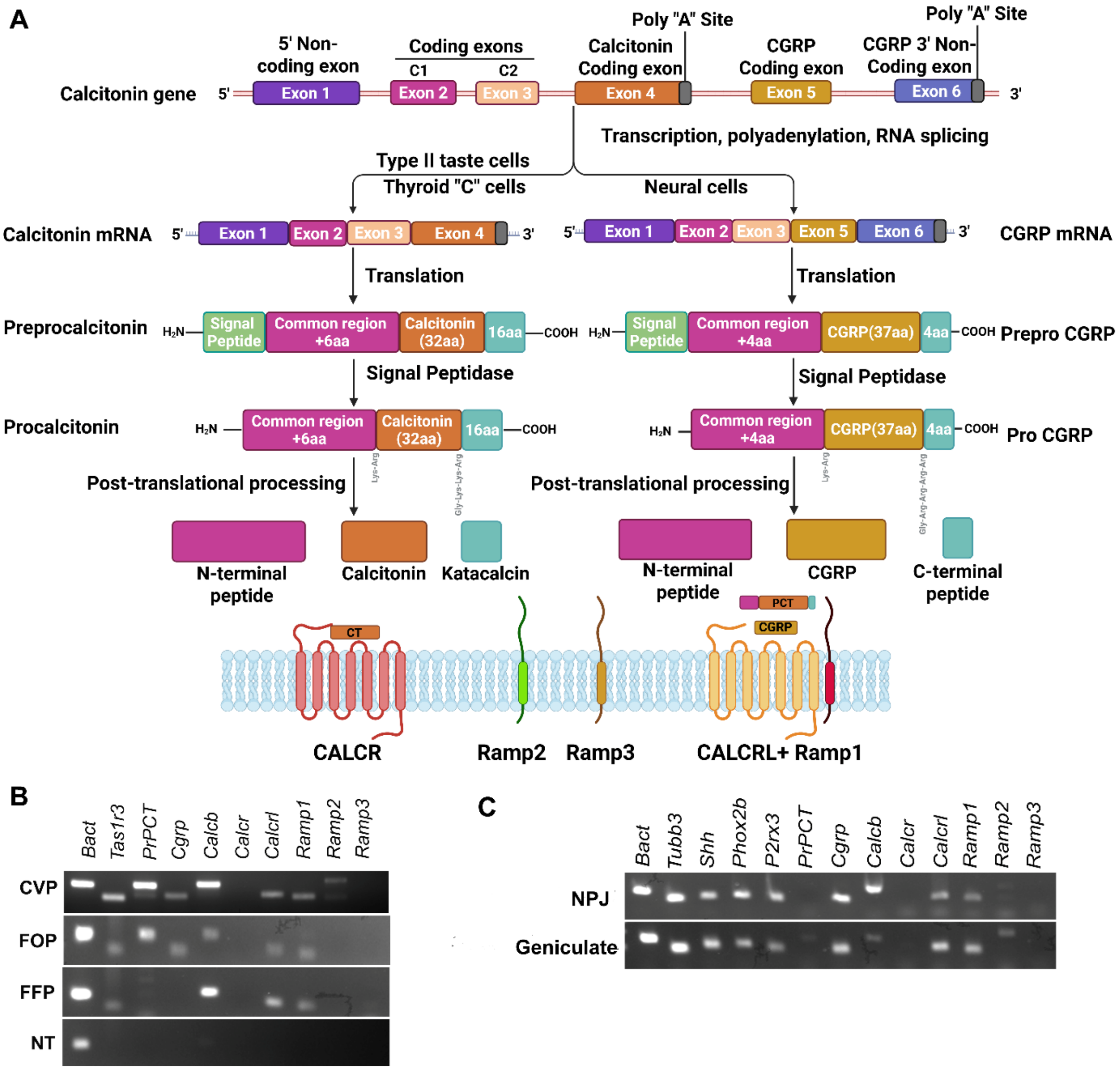


**Figure S1. *Calca* transcripts, peptides and their receptors, and their expression in taste papillae and ganglia**. **A)** The structure of the *Calca* gene, and the alternative splicing of the *Calca* mRNA that generates either the preprocalcitonin or pre proCGRP encoding transcripts. PCT, calcitonin (CT), Katacalcin and CGRP are produced by multistep proteolytic processing of the respective precursor proteins as shown. CT is a ligand for CALCR, while PCT and CGRP are ligands for CGRP1R, formed by a heterodimer of CALCRL and RAMP1. **B)** Expression of indicated transcripts in taste papillae and non-taste lingual epithelium (NT). *PrPCT, Calcb*, *Calcrl* and *Ramp1* expression is observed in CVP, FFP and

FOP. Weak *Cgrp expression* is observed in CVP and FOP, while it is undetectable in FOP and FFP. *Calcrl and Ramp1* are expressed in all taste papillae while *Calcr, Ramp2 and Ramp3 are* not detected in any tissue. None of the tested mRNAs are detected in NT. *Tas1r3* is used as a control to demonstrate the quality of taste cDNA, and *Bact* is used as cDNA synthesis control. **C)** qPCR of above transcripts in the geniculate and NPJ ganglia. *Cgrp*, *Calcb, Calcrl, Ramp1 and Ramp2* are expressed in both ganglia. Expression of *Ramp3*, *PrPCT* and *Calcr* is not observed in either ganglion. *Bact* is used as cDNA synthesis control, while the taste ganglion marker genes *Tubb3, Shh, Phox2b*, and *P2rx3* are used to demonstrate the quality of ganglia cDNA. Illustration in panel A was created in BioRender. Siddiqui, A. H. (2026) https://BioRender.com/gh3z1im

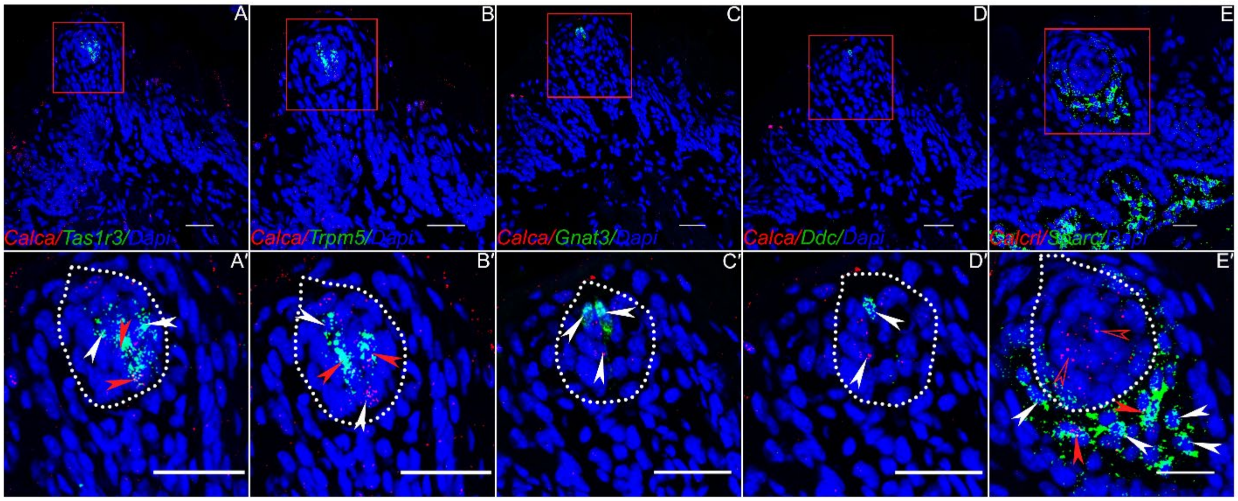


**Figure S2. Expression of *Calca* and *Calcrl* in fungiform papillae.** RNAscope Hiplex fluorescence assay was used to determine the coexpression of *Calca* with the taste cell markers *Tas1r3* (A-A'), *Trpm5* (B-B'), *Gnat3* (C-C'), *Ddc* (D-D') and of *Calcrl* with the fibroblast marker gene *Sparc*. The areas highlighted in red boxes in the top row are magnified in the bottom rows for each set. Taste buds are highlighted by white dotted lines. Areas inside white boxes in A-E are magnified in A'-E'. Weak expression of *Calca* is observed over all, and it appears to be coexpressed with *Tas1r3* and *Trpm5*. No coexpression is observed with *Gnat3* and *Ddc*. *Calcrl* expression is observed in basal (presumably stem/progenitor) cells in the taste buds and adjacent fibroblasts. Filled red arrowheads highlight double positive cells, and filled white arrowheads highlight single positive cells for taste/fibroblast marker genes, and open red arrowheads show single *Calcrl* positive cells. Scale bars = 30 μm. DAPI is used as a counterstain for nuclei.

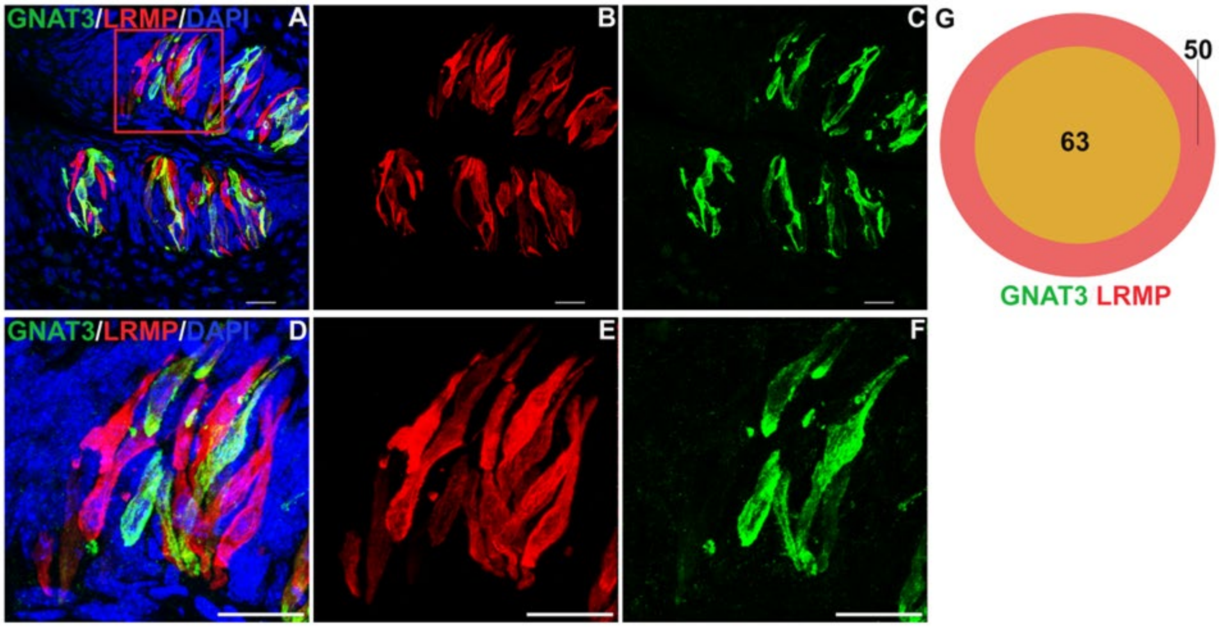


**Figure S3. Double labeling for GNAT3 and LRMP. A-F)** Unlike PCT staining shown in Figure 3, GNAT3 staining is restricted primarily to weaker LRMP expressing cells with less intense staining for LRMP. **G)** Venn diagram shows quantification of coexpression data. Scale bars= 30 µm. DAPI is used as a counterstain for nuclei.

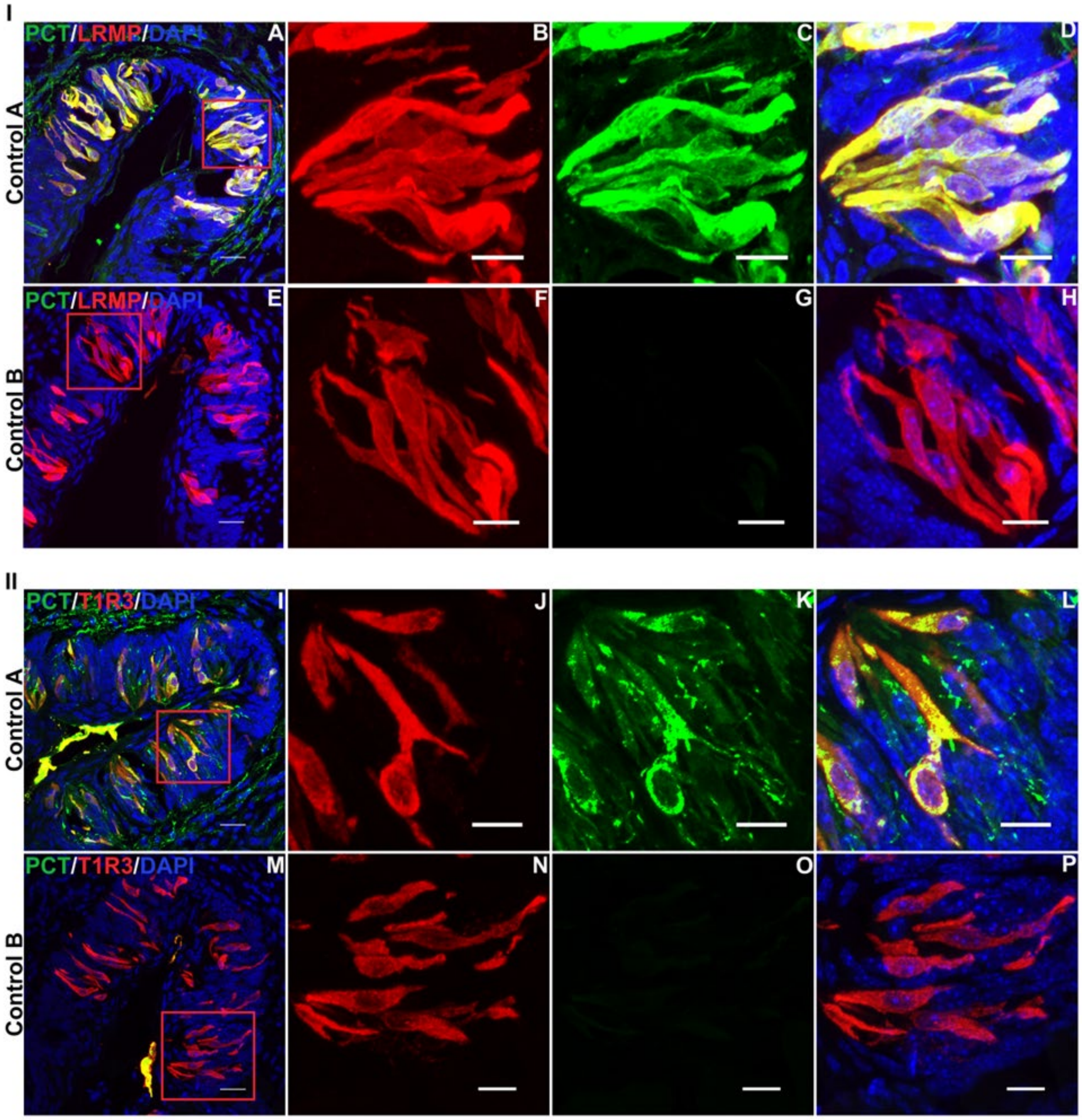


**Figure S4. Controls for immunostaining with two rabbit primary antibodies.** Double indirect immunofluorescence confocal microscopy of CVP shows cross-labeling when unlabelled donkey Fab fragment is omitted (control A, I & II), whereas adequate blocking of rabbit IgG is achieved with excess donkey Fab fragment (control B, I and II). Scalebars=30 µm. DAPI is used as a counterstain for nuclei.

| Probe | CAT number | LOT number |
|---|---|---|
| RNAscope™ HiPlex Probe- Mm-Ddc-T1 | 318681-T1 | 22363A |
| RNAscope™ HiPlex Probe- Mm-Tas1r3-T2 | 515431-T2 | 22174B |
| RNAscope™ HiPlex Probe- Mm-Sparc-T4 | 466781-T4 | 24017A |
| RNAscope™ HiPlex Probe- Mm-Calcrl-T5 | 452281-T5 | 24017A |
| RNAscope™ HiPlex Probe- Mm-Calca-T6 | 578771-T6 | 223634 |
| RNAscope™ HiPlex Probe- Mm-Gnat3-T8 | 531661-T8 | 22171A |
| RNAscope™ HiPlex Probe- Mm-Trpm5-T9 | 453251-T9 | 22363A |
| RNAscope™ HiPlex Probe- Mm-Arg1-T9 | 403441-T9 | 24017A |
| RNAscope™ HiPlex Probe- Mm-Ccr6-T11 | 424461-T11 | 24025A |

**Table S1:** List of RNAscope HiPlex Probes used in this study.

| Antibody | Host Species | Catalog No. | RRID | Source | Dilutions |
|---|---|---|---|---|---|
| PCT | Rabbit | LS-C296040 | - | LS Bio, Newark, CA | 1: 50 |
| GNAT3 | Goat | OAEB00418 | AB_1088282 | Aviva Systems Biology, San Diego, CA | 1:500 |
| LRMP | Rabbit | ORB166443 | - | Biorbyt, Durham, NC | 1:600 |
| T1R3 | Rabbit | - | - | Gift from Dr. Ichiro Matsumoto, Monell Chemical Senses center | 1:500 |
| CGRP | Goat | AB36001 | AB_725807 | Abcam Inc., Waltham, MA | 1:250 |
| TUBB3 | Rabbit | AB18207 | AB_444319 | Abcam Inc., Waltham, MA | 1:500 |

| | | | | | |
|---|---|---|---|---|---|
| Alexa Fluor 488 Conjugated AffiniPure Fab Fragmant Goat anti-Rabbit IgG (H+L) | Goat | 111-547-003 | - | Jackson Immuno Research Inc. West Grove, PA | 1:500 |
| Alexa Fluor 647 Conjugated AffiniPure Fab Fragmant Donkey anti-Rabbit IgG (H+L) | Donkey | 711-607-003 | - | Jackson Immuno Research Inc. West Grove, PA | 1:500 |
| Alexa Fluor 647 Conjugated AffiniPure Donkey anti-Goat IgG (H+L) | Donkey | 705-606-147 | - | Jackson Immuno Research Inc. West Grove, PA | 1:500 |
| Alexa Fluor 488 Conjugated Donkey anti-Goat IgG | Donkey | A11055 | - | Jackson Immuno Research Inc. West Grove, PA | 1:500 |
| AffiniPure Fab Fragment Donkey anti-Rabbit IgG(H+L) | Donkey | 711-007-003 | - | Jackson Immuno Research Inc. West Grove, PA | 1:500 |

**Table S2:** Primary and secondary antibodies for IHC and western blot used in this study.

| Gene | Forward Primer | Reverse Primer |
|---|---|---|
| *Bact* | GGCTGTATTCCCCTCCACG | CCAGTTGGTAACAATGCCATGT |
| *Calcrl* | CATCGTGGTGGCTGTGTTT | GTAATACAAGCTTCTGGCAATGG |
| *Calcr* | GCTGAGTGCAGAAACCCACT | TTTGCCTCATCTTGGTCACA |
| *Ramp1* | AGCCGCTTCAAGGAGAACAT | CGTGCTTGGTGCAGTAAGTG |
| *Ramp2* | TGAGGACAGCCTTGTGTCAA | CAGCACAGCAGAAAGGTTCC |
| *Ramp3* | AAGTTGGTTTTGGACGGTGA | GCATACCTGGGCACACTCA |
| *Tubb3* | GAACCTGGAACCATGGACAG | GTTGTTGCCAGCACCACTCT |
| *Shh* | AAGGATGAGGAAAACACGGG | GGTCACTCGCAGCTTCACTC |
| *Phox2b* | CACTTTTGGGGCCACGTC | CGTGGTCGGTGAAGAGTTTG |
| *P2rx3* | CCAAATATTCCTTCACTCGGC | GCTGCCATTCTCCATCTTGT |
| *PrePCT* | GGAGCAGGAGGAAGAGCAGG | GCCAGGTGCTTCAACCCCAA |
| *Cgrp* | TATGCAGATGAAAGCCAGGG | GTGGCAGTGTTGCAGGATCT |
| *Calcb* | CAGGAAGCTGGAACAGGAGG | AAGGCTTCAGAGCCCACATC |
| *Tas1r3* | CAAGTTCTTCAGCTTCTTCC | GGCGGCCACCCAGTTCCAGC |

**Table S3:** List of primers used for PCR and qPCR.
